## Supplemental Figures and Legends for "Ribonucleoprotein condensation driven by retrotransposon LINE-1 sustains RNA integrity and translation in mouse spermatocytes"

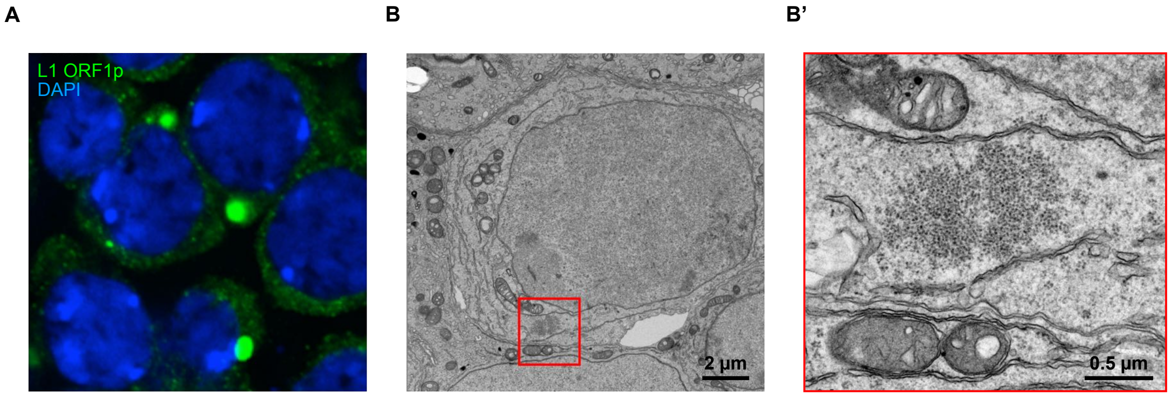


**Supplementary Figure 1.**

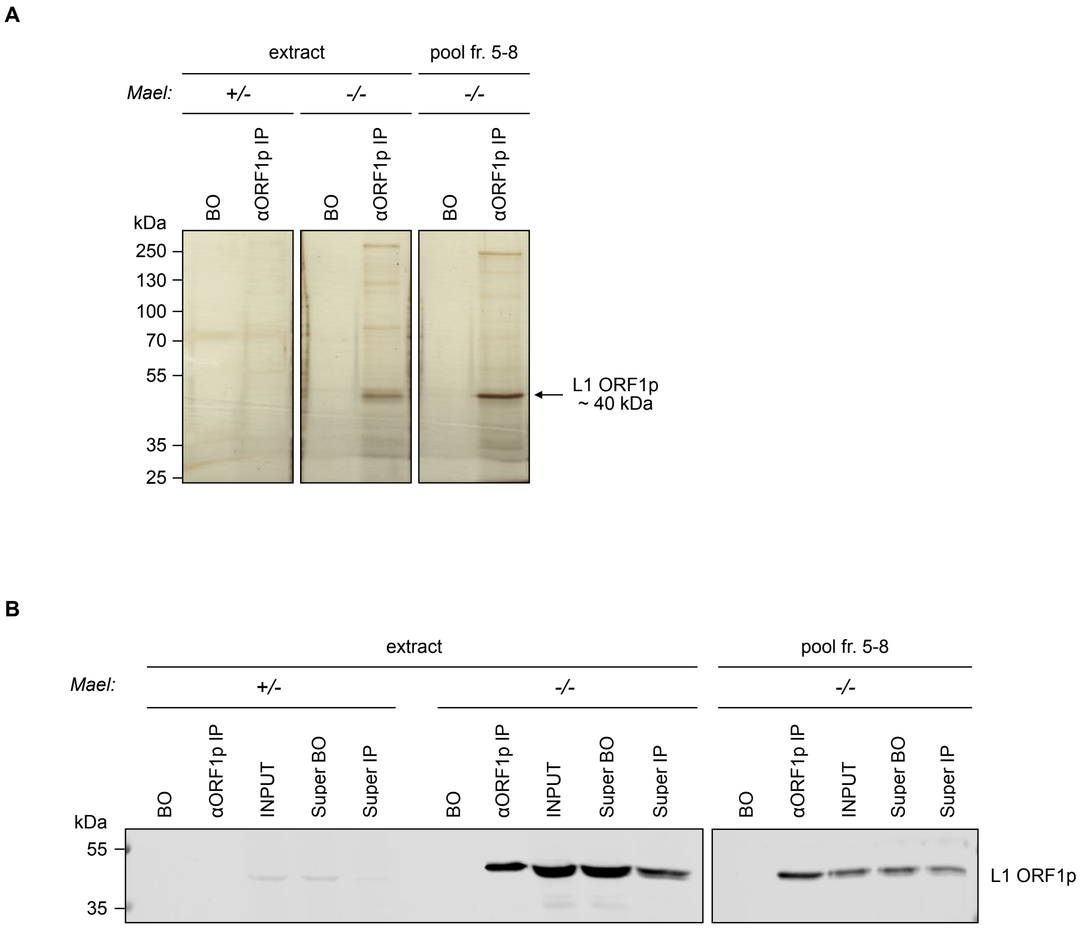


**Supplementary Figure 2.**

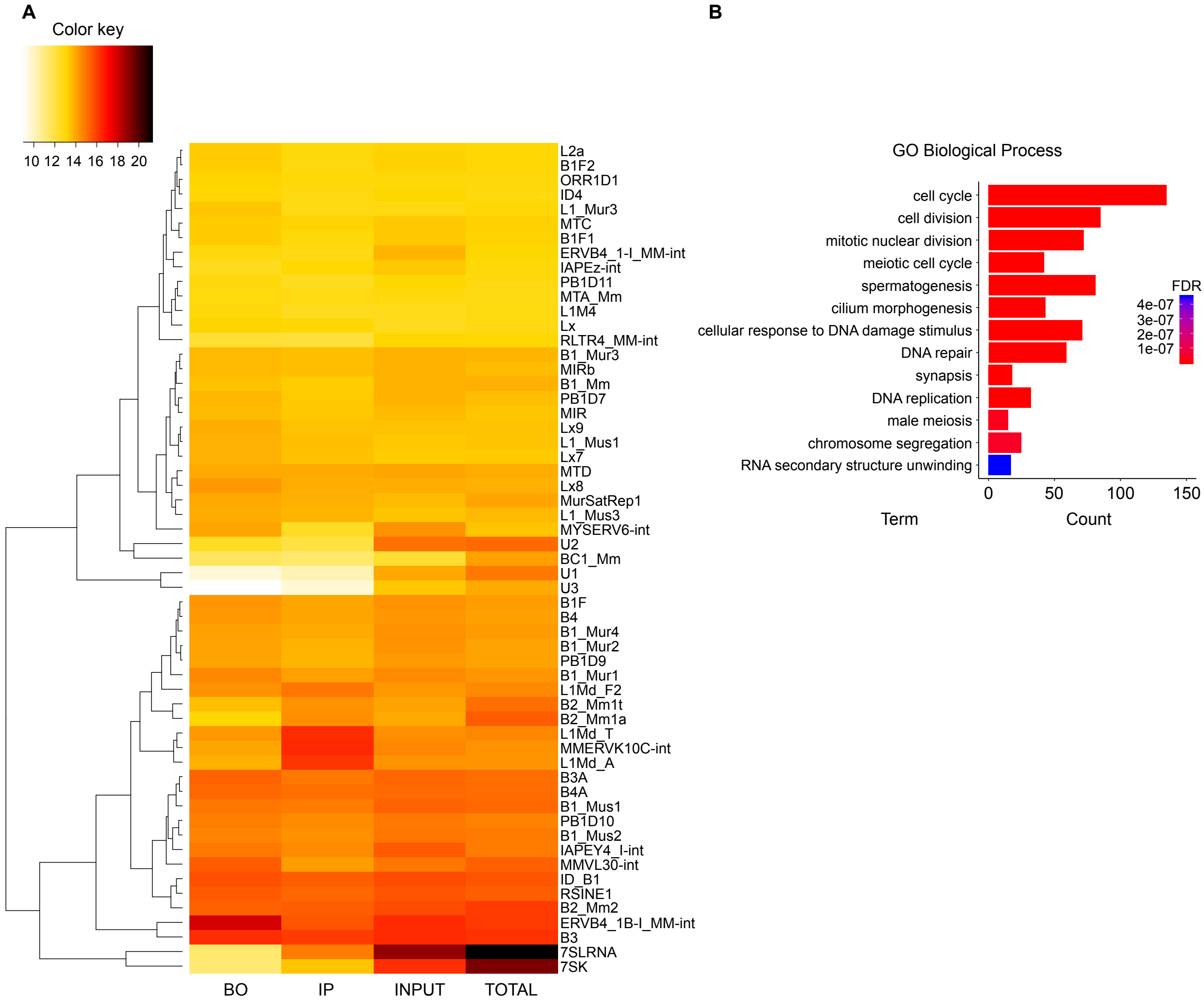


**Supplementary Figure 3.**

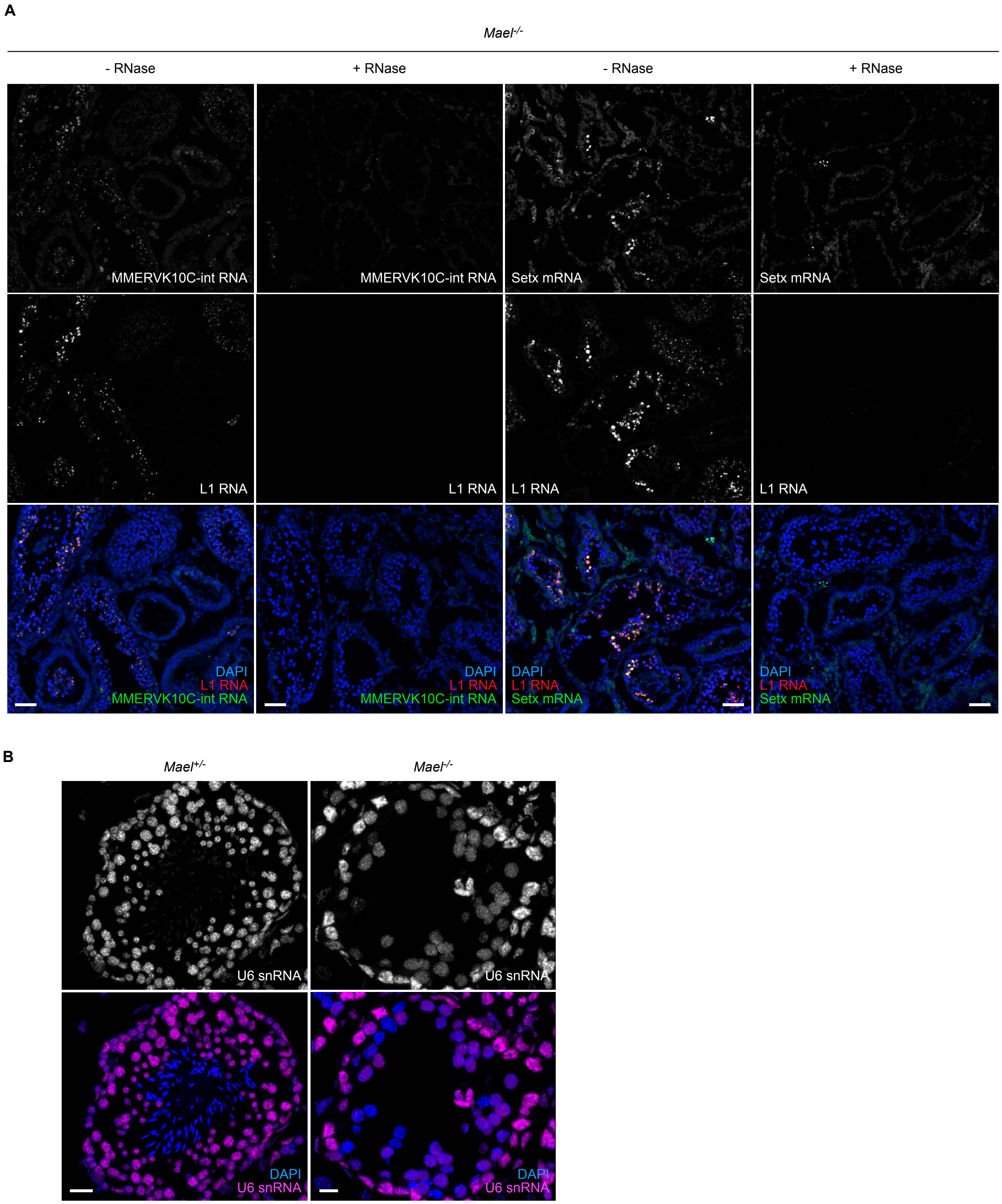


**Supplementary Figure 4.**

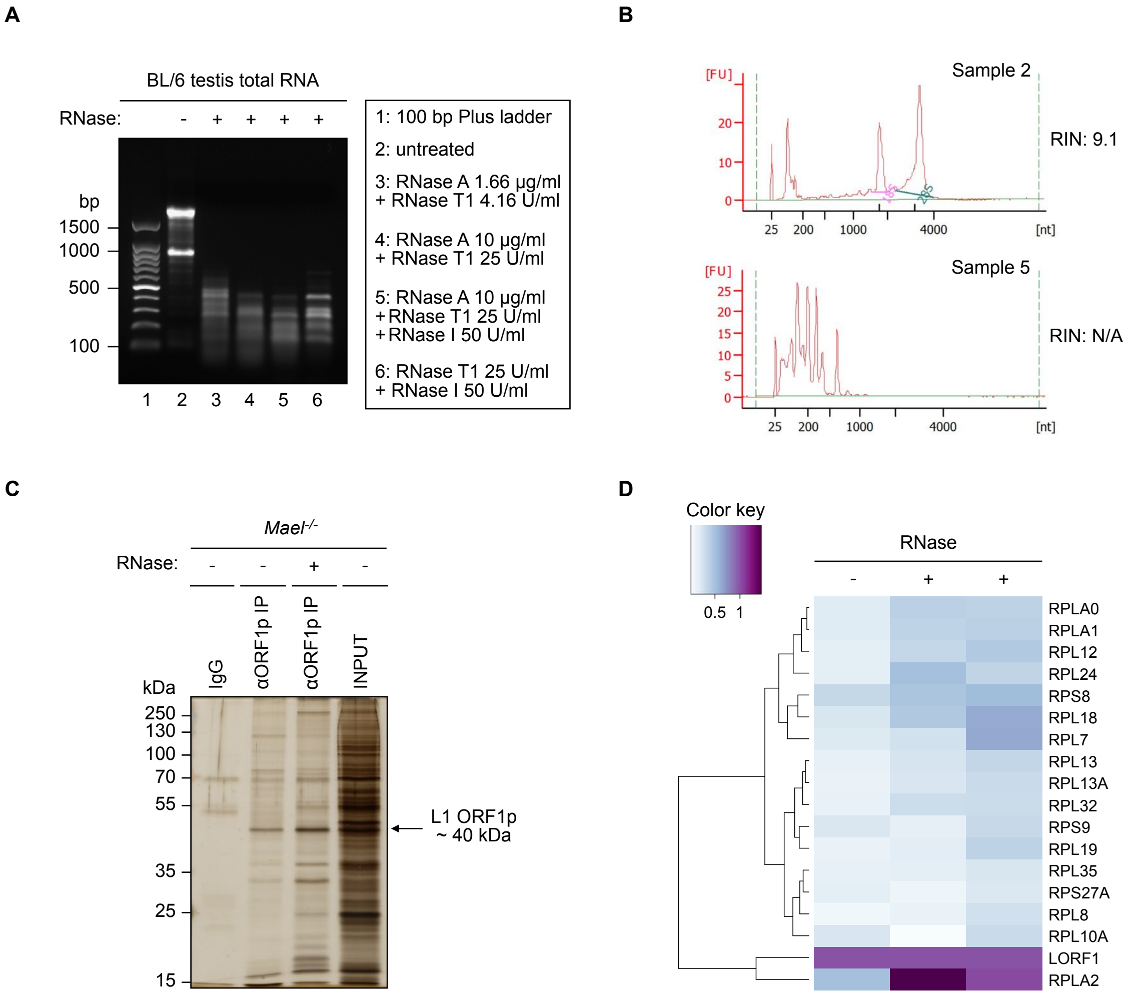


**Supplementary Figure 5.**

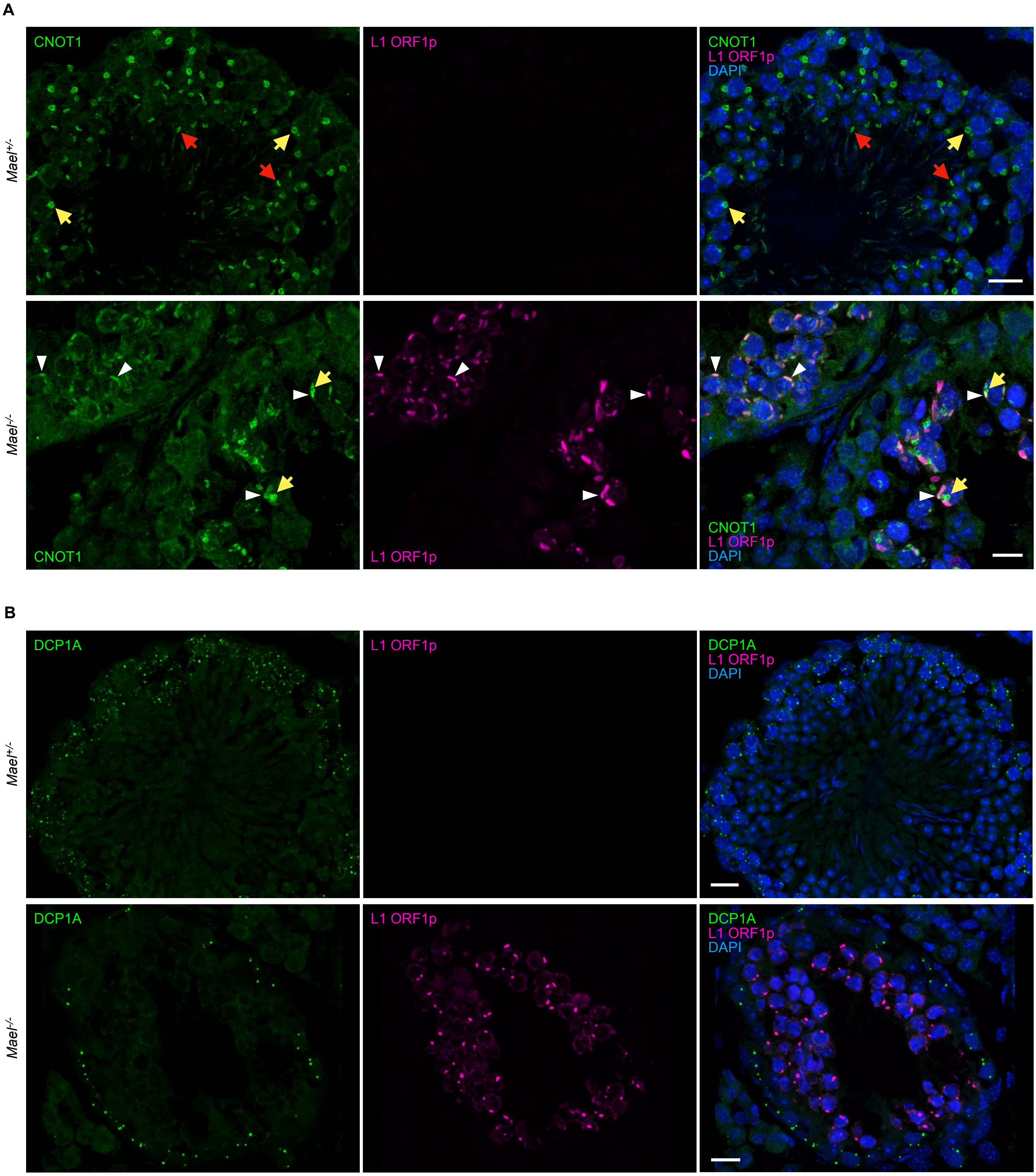


**Supplementary Figure 6.**

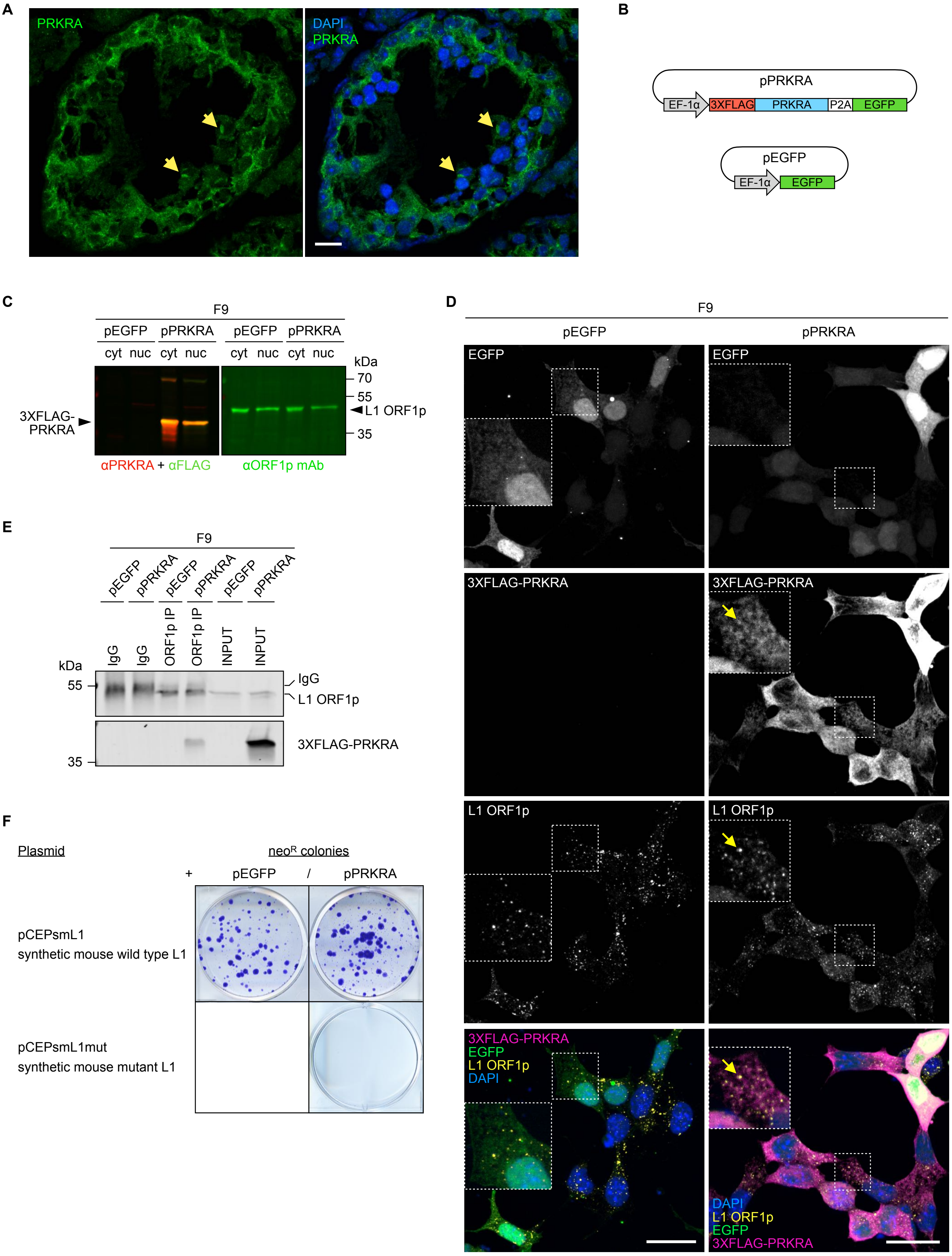


**Supplementary Figure 7.**

**Supplemental Figure S1. L1 ORF1p aggregates can be detected in the cytoplasm of spermatocytes from BALB/c wild-type mice. (A)** Immunofluorescence staining of L1 ORF1p (green) on BALB/c wild-type spermatocytes showing ORF1p accumulation in small cytoplasmic granules similar to early LBs. **(B – B')** Electron micrographs of a BALB/c wild-type spermatocyte harboring a small LB; boxed area in B is magnified in B'.
